## Supplemental Material for "Glycoproteomics of *Haloferax volcanii* reveals an extensive glycoproteome and concurrence of different *N*-glycosylation pathways"

**This document includes:**

Figures S1 to S5
Table S1

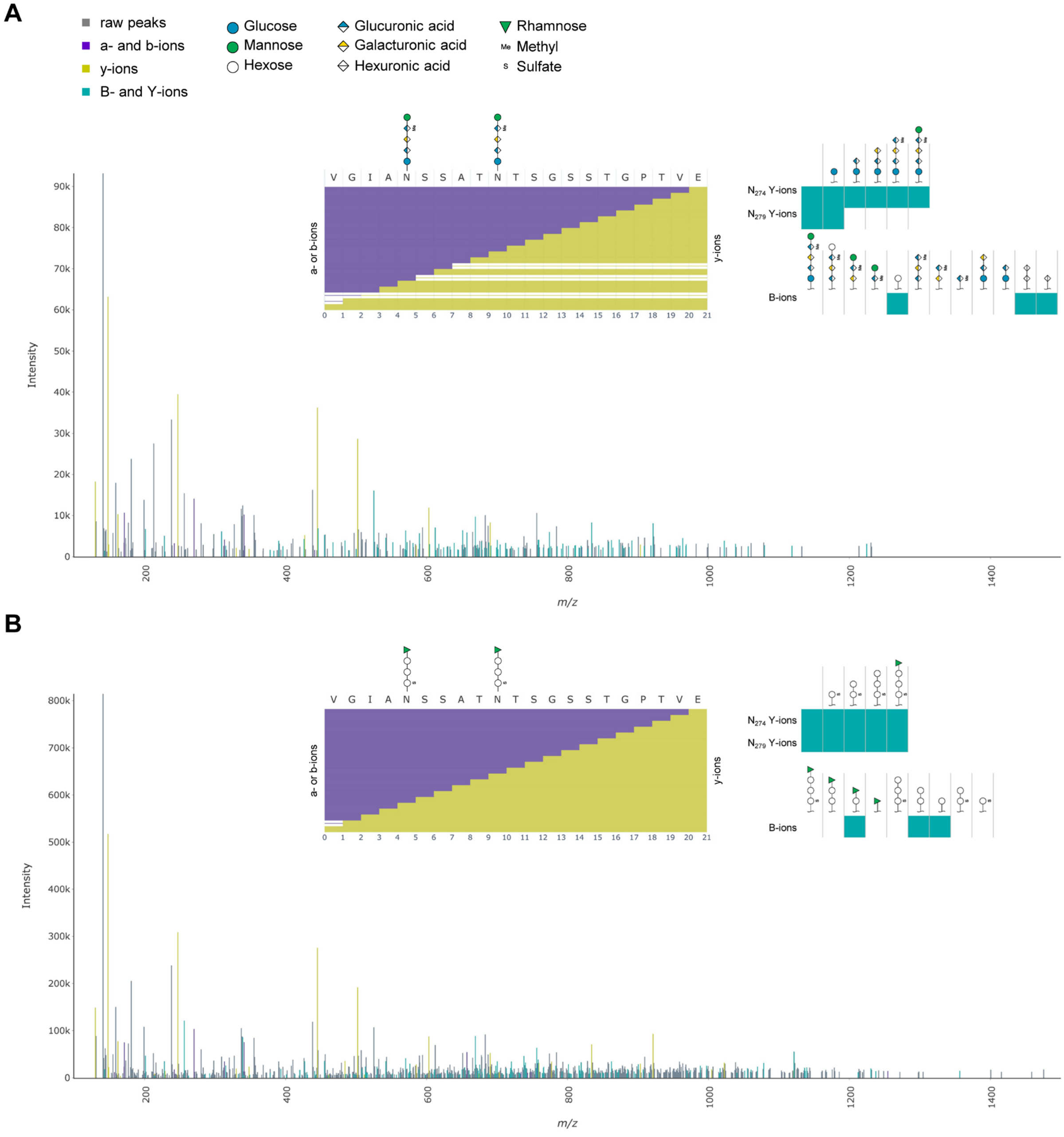


**Figure S1. MS2 spectra of SLG *N*-glycosites N274 and N279 strongly support modification by AglB- as well as Agl15-dependent *N*-glycans.** Annotated spectra for SLG *N*-glycopeptides comprising the peptide sequence VGIANSSATNTSGSSTGPTVE with AglB- (**A**) and Agl15-dependent (**B**) *N*-glycans attached to the *N*-glycosites N274 and N279. Measured raw peaks are shown in grey, annotated a- and b-ions in purple, y-ions in yellow and *N*-glycopeptide-specific Y- and B-ions in cyan. Insets illustrate the peptide sequence coverage through a- or b-ions (purple) and y-ions (yellow) (in both cases detected ions shown as wide bar, missing ions shown as line), as well as the coverage of Y- and B-ions (detected ions shown in cyan).

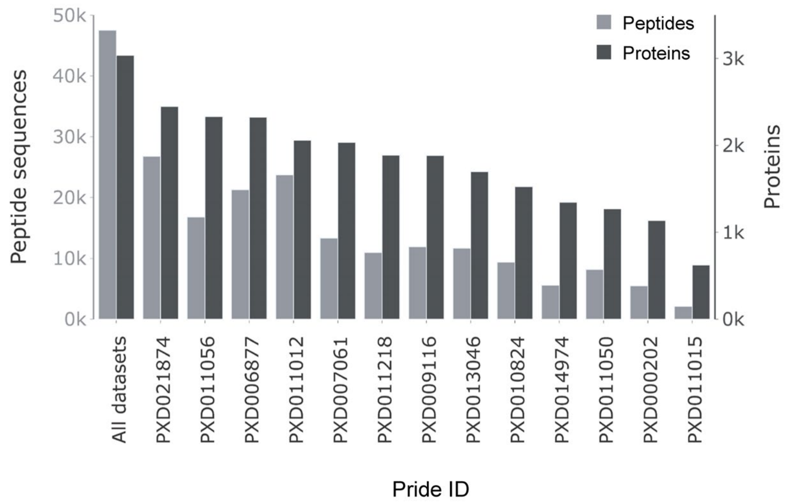


**Figure S2. More proteins and peptides have been identified in PXD021874 than in any other dataset of the ArcPP.** The number of identified peptides (light grey) and proteins (dark grey) for each dataset is shown as a barplot (sorted by the total number of identified proteins).

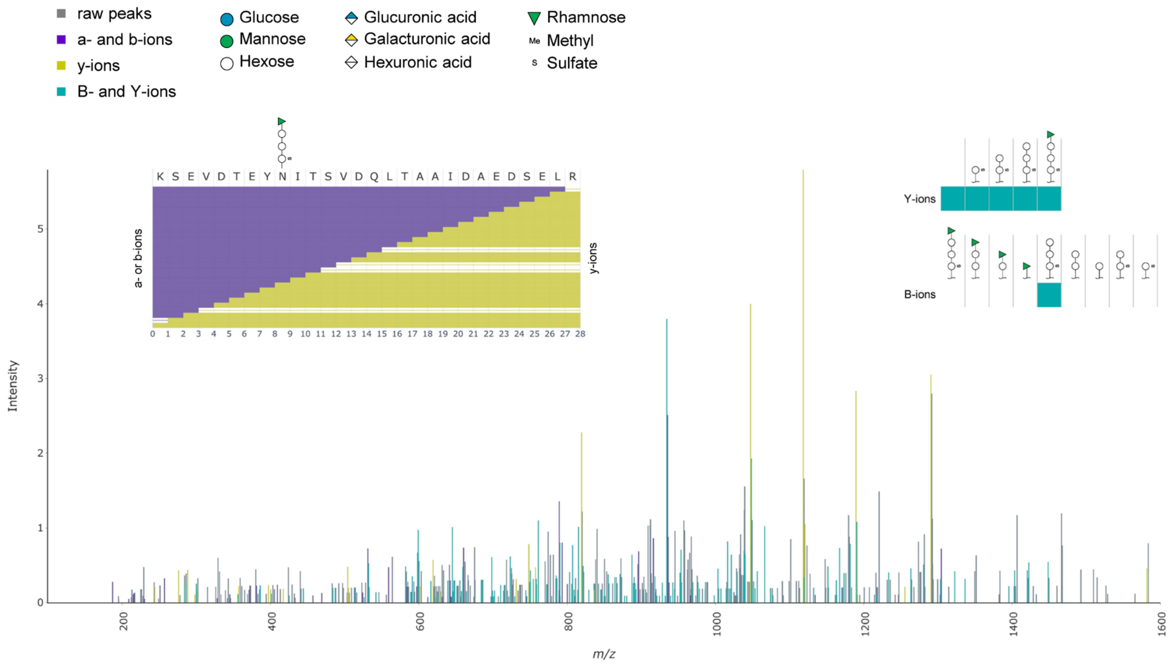


**Figure S3. Agl15-dependent *N*-glycosylation of Ths3 is strongly supported by MS2 fragment ion series.** Shown is an annotated spectrum for the Ths3 *N*-glycopeptide with sequence KSEVDTEYNITSVDQLTAAIDAEDSELR harboring an Agl15-dependent *N*-glycan. Measured raw peaks are shown in grey, annotated a- and b-ions in purple, y-ions in yellow and *N*-glycopeptide-specific Y- and B-ions in cyan. Insets illustrate the peptide sequence coverage through a- or b-ions (purple) and y-ions (yellow) (in both cases detected ions shown as wide bar, missing ions shown as line), as well as the coverage of Y- and B-ions (detected ions shown in cyan).

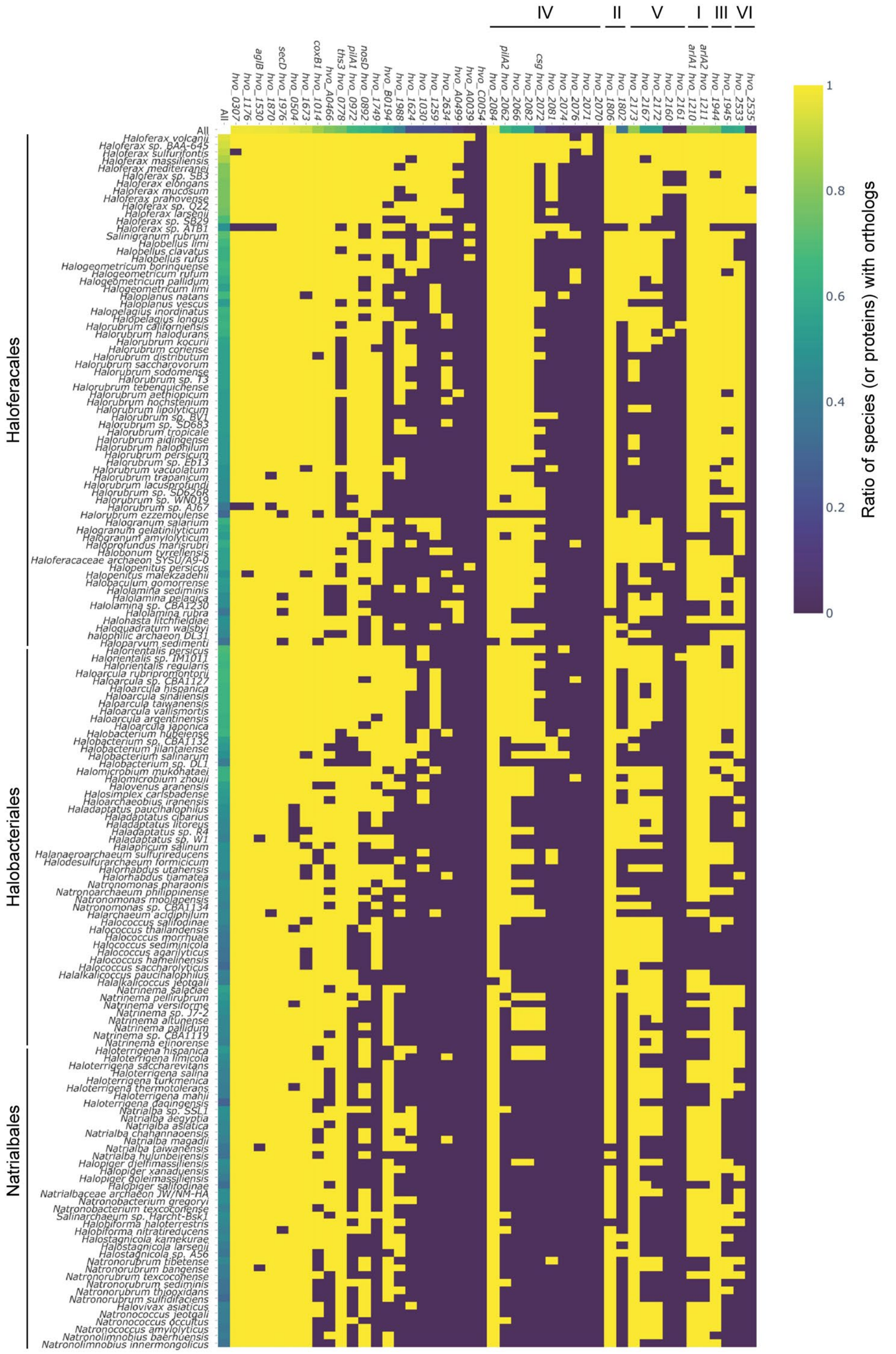


**Figure S4. Distribution of *N*-glycoproteins across Halobacteria.** For each identified *N*-glycoprotein (x-dimension) the percentage of species within the class of Halobacteria that carry an orthologue is presented. Furthermore, for each species (y-dimension), the percentage of *N*-glycoproteins for which an orthologue is present within the species' genome is shown. Percentages are shown as heat, ranging from yellow (100%) to purple (0%).

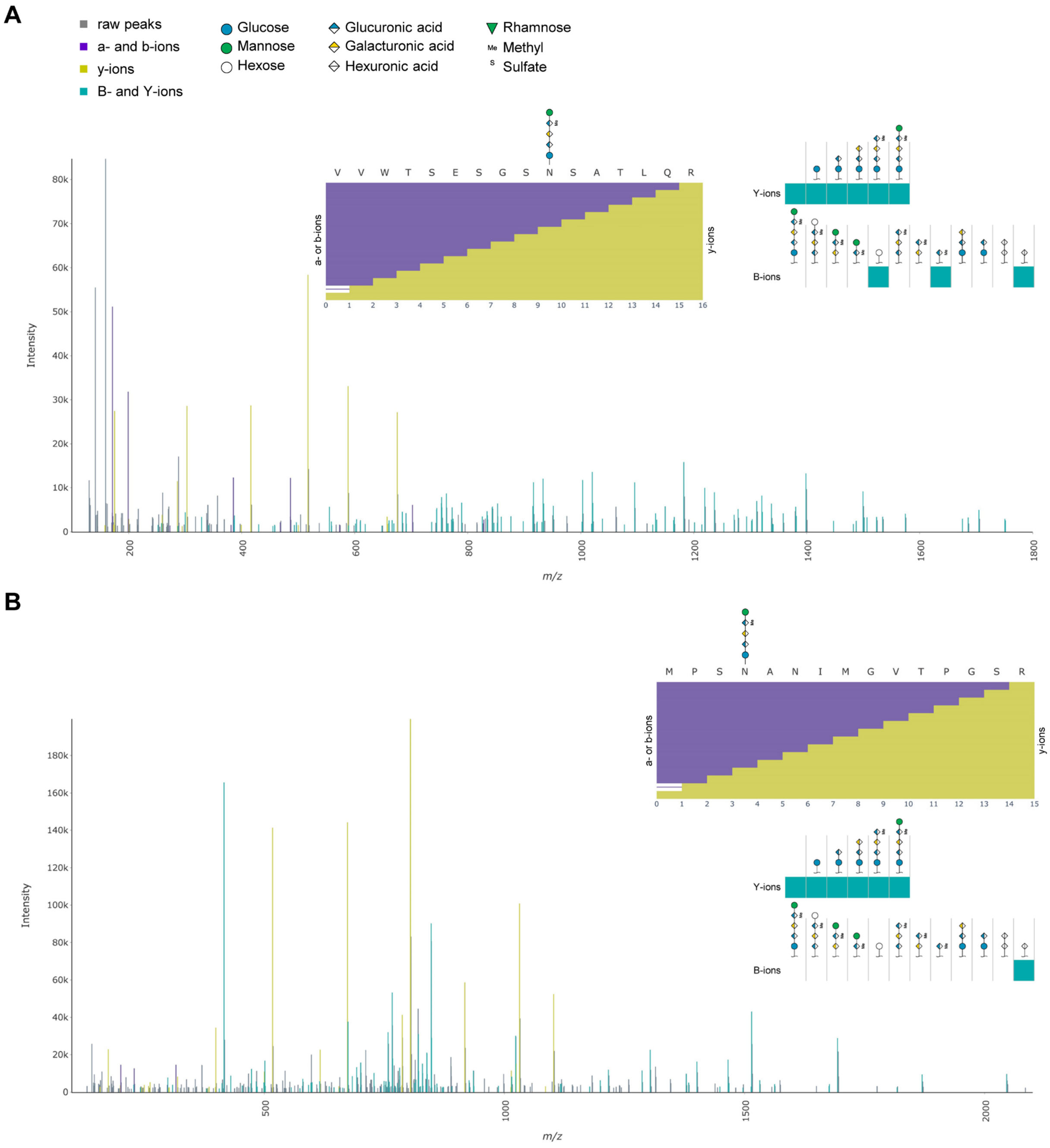


**Figure S5. *N*-glycopeptides with non-canonical *N*-glycosites are strongly supported by MS2 fragment ion series.** Annotated spectra for non-canonical *N*-glycosites within the peptide sequences VVWTSESGS**N**SATLQR (**A**) and MPS**N**ANIMGVTPGSR (**B**) corresponding to the protein pilA6 and HVO_2160, respectively, are shown. Both *N*-glycopeptides are modified by an AglB-dependent glycan and the site of attachment is indicated. Measured raw peaks are shown in grey, annotated a- and b-ions in purple, y-ions in yellow and *N*-glycopeptide-specific Y- and B-ions in cyan. Insets illustrate the peptide sequence coverage through a- or b-ions (purple) and y-ions (yellow) (in both cases detected ions shown as wide bar, missing ions shown as line), as well as the coverage of Y- and B-ions (detected ions shown in cyan).

**Table S1. Analysis of phylogeny, gene synteny and protein domains of identified *N*-glycoproteins.** For each protein that was identified to be *N*-glycosylated, the HVO ID, description, InterPro domains and arCOGlet classification is given together with results of phylogenetic and gene synteny analysis for the genes encoding the identified *N*-glycoproteins.

| HVO ID | Description[1] | InterPro Domains[2] | arCOG let[3] | Taxonomic range of orthologs[4] | Halobacteria with orthologs[5] | Neighboring genes/ SynTax analysis[2] | Comments |
|---|---|---|---|---|---|---|---|
| **HVO_0307** | conserved hypothetical protein | none | N | Hal+ | 157 | conserved gene clustering:<br>downstream:<br>HVO_0308 DUF1628 domain, pilin-related<br>HVO_0309 menG, menaquinone biosynthesis<br>upstream:<br>HVO_0306 probable transmembrane glycoprotein with HTH domain<br>HVO_0305/0304 etfA/B, subunits of an electron transfer flavoprotein | - |
| **HVO_0504** | DUF192 family protein | DUF192 | S | Arc | 152 | genomic vicinity of HVO_0505 to HVO_0511 in several genomes but no strict gene synteny:<br>downstream:<br>HVO_0505 small, uncharacterized<br>HVO_0506 ABC transporter permease/ATPase, unassigned substrate<br>HVO_0507 putative amidohydrolase, creatininase domain<br>HVO_0508 uncharacterized<br>HVO_0509 NUDIX family hydrolase<br>HVO_0510 mptD, folate biosynthesis<br>HVO_0511 azf, glucose-6-phosphate 1-dehydrogenase | tempting to speculate that HVO_0504 is a substrate-binding protein for HVO_0506 or that HVO_0506 transports the product of a potential enzymatic reaction catalyzed by HVO_0504 |
| **HVO_0778** | (Ths3) thermosome subunit 3 | - | O | Arc | 122 | - | - |
| **HVO_0892** | (NosD) ABC-type transport system periplasmic substrate-binding protein (probable substrate copper) | - | P | Hal+ | 102 | - | substrate assignment only tentative |
| **HVO_0972** | (PilA1) pilin PilA | DUF1628 | N | Eur | 107 | gene neighborhood not conserved | - |

| | | | | | | | |
|---|---|---|---|---|---|---|---|
| **HVO_1014** | (CoxB1) cox-type terminal oxidase subunit II | - | C | Arc | 136 | - | - |
| **HVO_1030** | DUF4382 domain protein | DUF4382 | S | Hal | 32 | gene neighborhood not conserved | - |
| **HVO_1176** | conserved hypothetical protein | none | S | Hal | 157 | part of a strictly conserved three-gene operon:<br>downstream:<br>HVO_1177 ADP-ribose pyrophosphatase<br>upstream:<br>HVO_1175 DUF2110 domain protein | - |
| **Cluster I** | | | | | | | |
| **HVO_1210** | (ArlA1) archaellin A1 | - | N | Eur | 131 | - | - |
| **HVO_1211** | (ArlA2) archaellin A2 | - | N | Eur | 131 | - | - |
| **HVO_1259** | conserved hypothetical protein | none | S | Hal | 32 | conserved gene clustering, 6-gene operon:<br>downstream:<br>HVO_1258 uncharacterized<br>HVO_1257 AAA-type ATPase, MoxR type<br>HVO_1256 vWFA domain<br>upstream:<br>HVO_1260 vWFA domain<br>HVO_1261 vWFA domain | - |
| **HVO_1530** | (AglB) dolichyl-monophosphooligosaccharide--protein glycotransferase AglB | - | M | Eur | 157 | - | - |
| **HVO_1624** | conserved hypothetical protein | none | S | Hal | 33 | one clustered gene:<br>upstream:<br>HVO_1622 ArsR family transcription regulator | - |
| **HVO_1673** | conserved hypothetical protein | none | S | Hal | 148 | gene neighborhood not conserved | - |
| **HVO_1749** | conserved hypothetical protein | none | S | Hal | 98 | gene neighborhood not conserved | - |
| **Cluster II** | | | | | | | |
| **HVO_1802** | peptidase M10 family protein | metalPep | E | Hal+ | 57 | - | the more specific assignment to metallopeptidase M10 is found for paralog HVO_1580 |

| | | | | | | | |
|---|---|---|---|---|---|---|---|
| **HVO_1806** | conserved hypothetical protein | none | S | Eur | 156 | gene neighborhood not conserved | - |
| **HVO_1870** | peptidase M50 family protein | pepM50 | O;R | Arc | 156 | three of five downstream genes are genomically clustered:<br>HVO_1871 hemQ coproheme decarboxylase<br>HVO_1874 oxidoreductase domain<br>HVO_1875 O-acetyltransferase domain | - |
| **<u>Cluster III</u>** | | | | | | | |
| **HVO_1944** | probable transmembrane glycoprotein / HTH domain protein | none | K | Hal+ | 124 | gene pairing conserved:<br>HVO_1944<br>HVO_1945 conserved hypothetical protein<br>no additional clustering | a HTH domain is assigned to close homolog C453_17564 |
| **HVO_1945** | conserved hypothetical protein | none | N | Hal | 92 | see HVO_1944 | - |
| **HVO_1976** | (SecD) protein-export membrane protein SecD | - | U | Eur | 156 | conserved gene pair:<br>HVO_1976 secD<br>HVO_1975 secF | - |
| **HVO_1988** | GATase domain protein | GATase | S | Hal | 68 | conserved gene pair:<br>HVO_1988<br>HVO_1987 sppA2 signal peptide peptidase<br>no additional clustering | - |
| **<u>Cluster IV</u>** | | | | | | | |
| **HVO_2062** | (PilA2) pilin PilA | DUF1628 | N | Eur | 107 | gene neighborhood not conserved | genes for the Agl15-dependent *N*-glycosylation pathway are encoded directly downstream of pilA2 (HVO_2046 to HVO_2061) |
| **HVO_2066** | conserved hypothetical protein | none | M | Hal+ | 95 | gene neighborhood not conserved | - |
| **HVO_2070** | conserved hypothetical protein | none | N | n/a | n/a | conserved gene pair:<br>HVO_2070<br>HVO_2069 RND family permease<br>no additional clustering | 40% protein sequence identity to HVO_A0466 |
| **HVO_2071** | probable secreted glycoprotein | none | M | Hal | 3 | clustering not analyzed (too rare) | - |
| **HVO_2072** | (Csg) S-layer glycoprotein | - | M | Hal | 36 | gene neighborhood not conserved | SLGs are highly diverse between species from the genus *Haloferax*; the SLG |

| | | | | | | | |
|---|---|---|---|---|---|---|---|
| | | | | | | | from *Hfx. gibbonsii* (24% protein sequence identity) has more orthologs (129) |
| **HVO_2074** | probable secreted glycoprotein | none | S | Hal | 10 | clustering not analyzed (too rare) | - |
| **HVO_2076** | probable secreted glycoprotein (nonfunctional) | DUF304 | M | Hal | 7 | clustering not analyzed (too rare) | targeted by transposon; most likely, a truncated version of this protein is stable enough to permit its proteomic detection; orthoDB analysis based on a close, nondisrupted homolog; DUF304 assigned to close homolog D320_03753 |
| **HVO_2081** | pectin lyase domain protein | pectin lyase | G;P | Hal | 19 | clustering not analyzed (too rare) | - |
| **HVO_2082** | conserved hypothetical protein | none | M | Hal+ | 95 | part of a three-gene operon:<br>HVO_2083 ABC transporter ATPase<br>HVO_2084 ABC transporter permease | tempting to speculate that HVO_2083/2084 transports the product of a potential enzymatic reaction catalyzed by HVO_2082; alternatively HVO_2082 could be a substrate-binding protein for HVO_2083/2084; this is considered less likely (lower number of orthologs; several genomes code for homologs of permease/ATPase but not for a homolog of HVO_2082) |
| **HVO_2084** | ABC-type transport system permease protein (probable substrate macrolides) | - | V | Arc | 157 | see HVO_2082 | - |
| **<u>Cluster V</u>** | | | | | | | |
| **HVO_2160** | probable secreted glycoprotein | Ig-like | M;O;<br>S | Eur | 9 | clustering not analyzed (too rare) | - |
| **HVO_2161** | probable secreted glycoprotein | none | T | Hal | 8 | clustering not analyzed (too rare) | - |

| | | | | | | | |
|---|---|---|---|---|---|---|---|
| **HVO_2167** | DUF4350 domain protein | DUF4350 GATase | S | Arc | 84 | conserved gene clustering (11 genes):<br><u>downstream:</u><br>HVO_2166 conserved hypothetical protein<br>HVO_2165 ABC transporter permease<br>HVO_2164 ABC transporter permease<br>HVO_2163 ABC transporter ATPase<br><u>upstream (total six genes):</u><br>HVO_2168 AAA-type ATPase, MoxR type<br>HVO_2169 conserved hypothetical protein<br>HVO_2170 conserved hypothetical protein<br>HVO_2171 DUF58 domain protein<br>HVO_2172 conserved hypothetical protein<br>HVO_2173 DUF1616 domain protein | - |
| **HVO_2172** | conserved hypothetical protein | none | L;M | Hal+ | 84 | see HVO_2167 | - |
| **HVO_2173** | DUF1616 family protein | DUF1616 | S | Eur | 133 | see HVO_2167 | - |
| **<u>Cluster VI</u>** | | | | | | | |
| **HVO_2533** | conserved hypothetical protein | none | S | Eur | 96 | gene neighborhood not conserved | - |
| **HVO_2535** | conserved hypothetical protein | none | S | Hal | 11 | clustering not analyzed (too rare) | - |
| **HVO_2634** | conserved hypothetical protein | none | S | Hal | 27 | gene neighborhood not conserved | - |
| **HVO_A0039** | conserved hypothetical protein | none | - | Hal | 5 | - | - |
| **HVO_A0466** | conserved hypothetical protein | none | N | Eur | 128 | conserved gene pair:<br>HVO_A0466<br>HVO_A0467 RND family permease<br>no additional clustering | 40% protein sequence identity to HVO_2070; SyntTax analysis performed with close homolog G3A49_05450 |
| **HVO_A0499** | conserved hypothetical protein | none | - | Hal | 12 | - | - |
| **HVO_B0194** | LppX domain protein | LppX | M | Eur | 86 | gene neighborhood not conserved | SyntTax analysis performed with close homolog G3A49_07715 |
| **HVO_C0054** | hypothetical protein | none | - | n/a | 0 | - | - |

[1]the description term “conserved hypothetical protein” is used for proteins devoid of an InterPro domain assignment and for which functionally characterized homologs are absent or too distant for annotation transfer.
[2]with few exceptions, InterPro domains, neighboring genes and SyntTax analyses are not reported if the *N*-glycoprotein itself is well characterized. For plasmid-encoded proteins (codes start with HVO_A, HVO_B, or HVO_C), SyntTax analysis is not available, unless a close homolog from a related species is chromosomally encoded. Domain abbreviations: DUF, domain of unknown function; GATase, glutamine aminotransferase; metalPep, metallopeptidase; pepM50, MEROPS peptidase family M50; vWFA, von Willebrand factor type A.
[3]arCOGlet: C, energy production and conversion; E, amino acid transport and metabolism; G, carbohydrate transport and metabolism; K, transcription; L, replication, recombination and repair; M, cell wall/membrane/envelope biogenesis; N, cell motility; O, post-translational modification, protein turnover, chaperones; P, inorganic ion transport and metabolism; R, general function prediction only; S, function unknown; T, signal transduction mechanisms; U, intracellular trafficking, secretion, and vesicular transport; V, defense mechanisms
[4]taxonomic range: Hal, Halobacteria; Eur, Euryarchaeota; Arc, Archaea; Hal+ indicates few additional orthologs beyond Halobacteria.
[5]number of genomes from the taxonomic order Halobacteria which contain an ortholog; a total of 161 genomes are considered. The term “n/a” refers to proteins which are not assigned to a protein group in OrthoDB.
